# A New Big-Data Paradigm for Target Identification and Drug Discovery

**DOI:** 10.1101/134973

**Authors:** Neel S. Madhukar, Prashant K. Khade, Linda Huang, Kaitlyn Gayvert, Giuseppe Galletti, Martin Stogniew, Joshua E. Allen, Paraskevi Giannakakou, Olivier Elemento

**Affiliations:** Institute for Computational Biomedicine, Dept. of Physiology and Biophysics, Weill Cornell Medical College, New York, NY 10065, USA; Institute for Precision Medicine, Weill Cornell Medical College, New York, NY 10065, USA; Meyer Cancer Center, Weill Cornell Medical College, New York, NY 10065, USA; Tri-Institutional Training Program in Computational Biology and Medicine, New York, NY 10065, USA; Division of Hematology and Medical Oncology, Department of Medicine, Weill Cornell Medical College, New York, NY 10065, USA; Oncoceutics, Inc., Philadelphia, PA 19104, USA

**Author notes:** Co-first authors (Neel S. Madhukar, Prashant K. Khade). Correspondence: Joshua Allen, Paraskevi Giannakakou, or Olivier Elemento.

## Abstract

Drug target identification is one of the most important aspects of pre-clinical development yet it is also among the most complex, labor-intensive, and costly. This represents a major issue, as lack of proper target identification can be detrimental in determining the clinical application of a bioactive small molecule. To improve target identification, we developed BANDIT, a novel paradigm that integrates multiple data types within a Bayesian machine-learning framework to predict the targets and mechanisms for small molecules with unprecedented accuracy and versatility. Using only public data BANDIT achieved an accuracy of approximately 90% over 2000 different small molecules – substantially better than any other published target identification platform. We applied BANDIT to a library of small molecules with no known targets and generated ∼4,000 novel molecule-target predictions. From this set we identified and experimentally validated a set of novel microtubule inhibitors, including three with activity on cancer cells resistant to clinically used anti-microtubule therapies. We next applied BANDIT to ONC201 – an active anti- cancer small molecule in clinical development – whose target has remained elusive since its discovery in 2009. BANDIT identified dopamine receptor 2 as the unexpected target of ONC201, a prediction that we experimentally validated. Not only does this open the door for clinical trials focused on target-based selection of patient populations, but it also represents a novel way to target GPCRs in cancer. Additionally, BANDIT identified previously undocumented connections between approved drugs with disparate indications, shedding light onto previously unexplained clinical observations and suggesting new uses of marketed drugs. Overall, BANDIT represents an efficient and highly accurate platform that can be used as a resource to accelerate drug discovery and direct the clinical application of small molecule therapeutics with improved precision.

## Introduction

It typically takes 15 years and 2.6 billion dollars to go from a small molecule in the lab to an approved drug ^1-3^, and for natural products and phenotypic screen derived small molecules, one of the greatest bottlenecks is identifying the targets of any candidate molecules^2,4^. Proper understanding of binding targets can position drugs for ideal indications and patients, allow for better analog design, and explain observed adverse events. There exist a number of experimental approaches for target identification ranging from affinity pull-downs to genome-wide knockdown screens ^4,5^, but these approaches are labor, resource, and time intensive, not to mention failure prone. Computational target prediction has the potential to substantially reduce the work and resources needed for drug target identification. Existing computational methods traditionally fall into three major categories: ligand-based, molecular docking, and data driven. Ligand-based approaches take known binding targets for a given drug and attempt to find other proteins that are sufficiently similar to the known targets ^6,7^. These similar proteins are then predicted as novel targets. However, to achieve high predictive power they require a large input of known binding partners for each tested drug, and therefore can only be used on drugs which have prior comprehensive target information ^6,7^. Because of this, these methods are often not broadly applicable, especially to orphan molecules – molecules with no known binding targets. On the other hand, molecular docking uses simulations of small molecules interacting with proteins to model if and how a drug may bind a given protein ^8,9^. However, this approach requires significant computational power and complex 3D structures for each queried protein – data that is often unavailable.

Traditionally, data-driven methods have focused on a single aspect out of a small molecule’s activity in a biological system. Wang et al. ^10^ used post-treatment gene expression changes to predict drugs with shared targets ^11,12^. Another method relied on side-effect similarity between drugs with known targets to predict new drug-protein interactions ^13^. However, this method was restricted to the small subset of small molecules that had been clinically tested and had thorough side effect annotation. Though each of these methods represents a significant advancement in the field, they all suffer from either lack of accuracy or broad utility – evidenced either by an inability to reliably validate target predictions, or by their limited applicability to a small subset of all small molecules. This is not very surprising though, as past research has demonstrated that these individual datasets are noisy, thus, it is expected that reliance on any single data type will lead to low predictive power ^14-16^.

Additionally, other groups have shown how the combination of multiple types of data can improve the calculation of drug-drug similarities^17^ and adverse event prediction^18^, yet, this type of combinatorial approach has not been fully explored for drug-target prediction. The few reported studies using combinatorial approaches for drug-target prediction, suffer from significant limitations that minimize their impact in the field. These limitations include the use of gene-based similarity features, a method inherently biased against the discovery of diverse types of targets (favoring instead, the discovery of genes of the same class as the known drug-targets), the small number of drugs used in the study (<500), or lack of experimental target validation^19-21^. To overcome these limitations, we introduce BANDIT, a novel drug-target prediction platform. BANDIT achieves unprecedented target-identification accuracy, without any reliance on gene-based similarities (making it broadly applicable to newly discovered compounds), uncovers novel targets for the treatment of cancer, and can be used to quickly pinpoint potential therapeutics with novel mechanisms of action to accelerate drug development.

### A novel combinatorial Big-Data Approach leads to a large increase in predictive power

In the age of “Big Data” there has been an explosion of techniques that permit genomic, chemical, clinical, and pharmacological measurements to characterize a small molecule’s mechanism. Many such measurements are either already published or are reasonably straightforward to perform. We hypothesized that integrating the multiple, independent pieces of evidence provided by each data type into a cohesive prediction framework would dramatically improve target predictions. To test this hypothesis, we developed **BANDIT**: a **B**ayesian **AN**alysis to determine **D**rug **I**nteraction **T**argets. BANDIT integrates over 20,000,000 data points from six distinct data types – drug efficacies^22^, post-treatment transcriptional responses ^11,12^, drug structures ^23,24^, reported adverse effects ^25^, bioassay results ^23,24^, and known targets ^26,27^ – to predict drug-target interactions. This underlying database contains information on approximately 2,000 different drugs with 1,670 different known targets and over 50,000 unique orphan compounds (compounds with no known targets).

For each data type we calculate a similarity score for all drug pairs with known targets. Since each dataset uses a distinct reporting metric, the similarity calculation was specific to the data type being considered (**Figure S1; Methods**). Previous approaches have argued that high similarity in one feature indicates high similarity in others, implying that only one or two data types are sufficient for target prediction since others can be inferred ^28^. However, using our vastly expanded dataset, we found little overall correlation across different similarity scores (Figure 1A; **Figure S2**). These results suggest that each data type is measuring a distinct aspect of a molecule’s activity and that individual features for a given drug cannot be extrapolated based on other data types. This shortcoming further supported our hypothesis that a novel approach that integrates independent data types could significantly improve target prediction accuracy.

**Figure 1:**
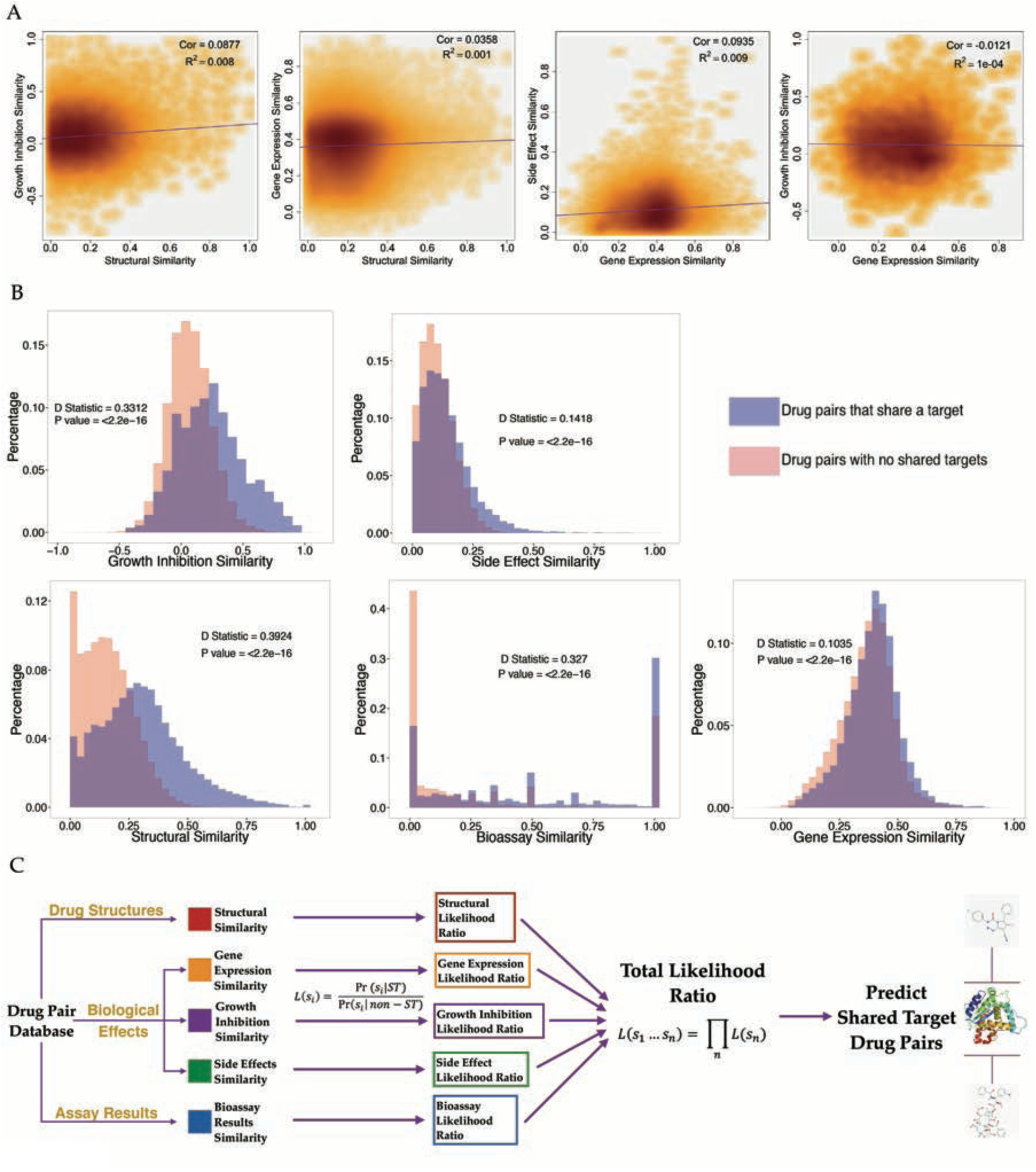
BANDIT exploits both the independence and individual predictive powers of each data type – A) Density plots showing how various different similarity scores correlate with one another, with darker area corresponding to a higher density of values. R^2^ and P value were calculated using a pearson correlation. B) Distributions of similarity scores across two sets – drug pairs known to share a target and those with no known shared targets. P values and D statistics were calculated using the Kolmogorov-Smirnov test. C) Schematic of BANDIT’s method of integrating multiple data types to predict shared target drug pairs.

We next separated drug pairs into those that shared at least one known target (>34,000 pairs) and pairs with no known shared targets (>1,250,000 pairs). We applied a Kolmogorov-Smirnov test to each similarity score and used the associated D statistic to calculate the degree a given data type could separate out drug pairs that shared targets (Figure 1B). We found that all features were able to significantly separate the two classes (P < 2e-16), and structural similarity was found to be the most discriminative among all features evaluated (D_Structure_ = 0.39). Additionally, we discovered that similarity across an unbiased set of bioassays and the relatively simple NCI-60 growth inhibition screen could strongly differentiate shared target drug pairs (D_Bioassay_ = 0.327 & D_GI50_ = 331), while, surprisingly ^10,13,29^, transcriptional responses (D_TResponse_ = 0.1) and reported adverse effects (D_SideEffect_ = 0.14) were much weaker differentiators. This information not only identifies the strengths of each data type, but will also allow researchers to efficiently prioritize experiments when faced with limited resources.

For every drug pair, BANDIT converts each individual similarity score into a distinct likelihood ratio. These individual likelihood ratios are then combined within a Naïve Bayes framework to obtain a total likelihood ratio (TLR) that is proportional to the odds of two drugs sharing a target given all available evidence (Figure 1C; **Methods**). We calculated TLRs for all possible drug pairs with known targets and the output was evaluated using 5-fold cross validation. We observed an Area Under the Receiver Operating Curve (AUROC) of 0.89 –higher than any competing approach ^13,28^– demonstrating that BANDIT’s integrative approach can accurately identify drugs that share targets. We recomputed the AUROC while varying the number of included data types and observed an overall increase in predictive power as we added new data types (Figure 2A). Furthermore we observed a steady increase in predictive power regardless of the addition order. This result verified the power of BANDIT’s “Big Data” approach and demonstrated how separate information sources can be combined to yield predictions more powerful than those obtained from any individual source **(Figure S3)**. This was confirmed using the KS test where we saw that the TLR output could better separate shared target drug pairs than any individual similarity score with a drastic increase in performance when focusing on drug pairs with all 5 data types (D_TLR_ = .69, **Figure S4**). Furthermore, we observed that BANDIT’s ratio of true to false positives continually increased as we raised the cutoff value, indicating that BANDIT’s TLR output is a dynamic value that estimates the strength and confidence level of a specific prediction and can effectively pick out high quality shared-target predictions (Figure 2B, **Figure S5**).

**Figure 2:**
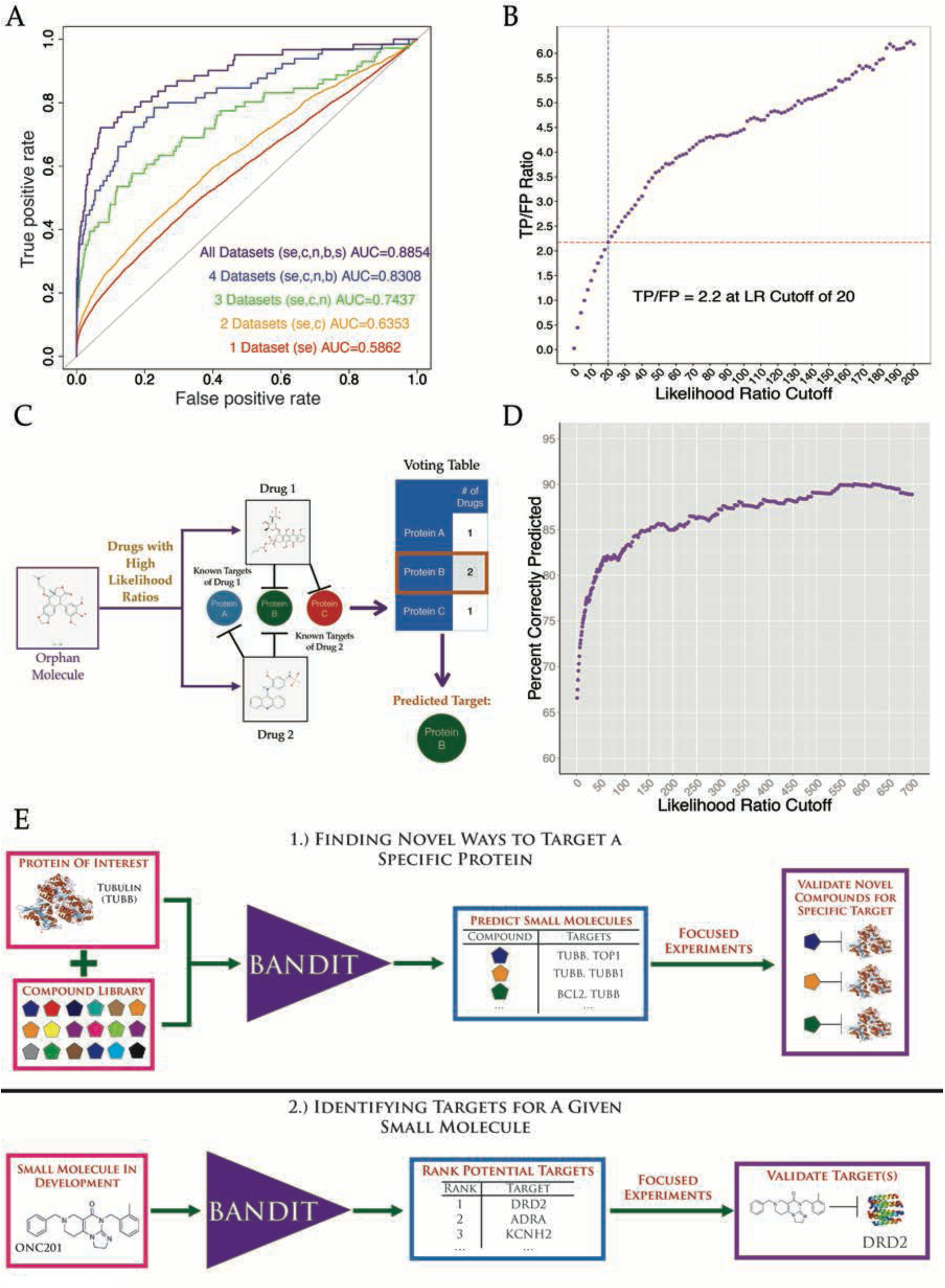
BANDIT can accurately predict shared targets and specific target interactions – A) Area under the receiver-operating curve for different sets of data types. SE = Side effects; C = CMap; N = NCI60; B = Bioassays; S = Structure. B) Ratio of true positives to false positives at different likelihood ratio cutoffs. C) Schematic of the BANDIT voting schematic for predicting specific target interactions. D) Accuracy level of BANDIT’s voting algorithm at various likelihood ratio cutoffs E) Schematic of two proposed operating scenarios for BANDIT

### BANDIT can replicate the results of experimental screens and predict specific target interactions

We next investigated how we could use BANDIT to replicate results from published experimental screens. Peterson et al. ^30^ tested 178 known protein kinase inhibitors against a panel of 300 different kinases and measured the level of inhibition (in terms of percent remaining kinase activity) for each inhibitor-kinase pair. We examined all orphan molecules – molecules with no known targets – in both the Peterson kinase database and BANDIT’s, and, used BANDIT to predict potential kinases targets for each orphan molecule (**Methods**). We observed that the kinase targets BANDIT predicted for each orphan molecule had higher levels of reported inhibition in the Peterson dataset than non-predictions (p<1e-5; **Figure S6**). This result supports using BANDIT to guide experimental screens while minimizing operational costs.

Moving forward from shared-target predictions, we examined whether for a given drug BANDIT could be used to predict a specific binding target from our database of over 1,600 unique proteins. We hypothesized that if a protein appeared as a known target in a large number of shared target predictions, then it is likely a target for the tested orphan molecule. To test this hypothesis, we developed a “voting” algorithm to predict specific targets for each orphan small molecule by identifying any recurring targets (Figure 2C, **Methods**). We applied our voting method to all drugs in our database with known targets and demonstrated that as we required more stringent TLR values for a pair of drugs to be predicted to share a target, the accuracy level – measured by whether BANDIT correctly identified a known drug target – steadily increased (Figure 2D). The accuracy level eventually reached ∼90%, demonstrating that BANDIT could be used to accurately identify specific targets for a diverse set of small molecules.

We then used BANDIT to predict novel targets for 14,168 small molecules with no known targets or mechanisms of action in our database. We confidently predicted targets for 4,167 unique small molecules (30% of our original set), with predictions spanning over 560 distinct protein targets. By setting a higher TLR cutoff for predictions and requiring a higher number of “votes” for any predicted targets, we further narrowed this list to 720 high confidence target predictions. To date, this is the largest database of novel drug-target predictions (nearly double the number of drugs in DrugBank’s drug-target database) and this list can be interrogated further to discover novel therapeutics and small molecules for a target of interest. Based on this success, we envisioned two main operating scenarios for BANDIT: 1) Using BANDIT in combination with the library of orphan small molecules to identify new small molecules targeting a specific protein and 2) to integrate BANDIT directly into the drug development pipeline to predict targets and guide experiments for drugs currently in development (Figure 2E).

### Discovery of Novel Microtubule-Targeting Compounds Capable of Overcoming Drug Resistance

Beginning with the first operating scenario, we used BANDIT to identify novel ways to target microtubules. Anti-microtubule drugs make up one of the largest and most widely used classes of cancer chemotherapeutics, with tubulin being one of the most validated anticancer targets to date ^31-34^. Interestingly, and unlike most classes of cancer chemotherapy drugs or targeted-therapies in oncology, microtubule inhibitors are further sub-categorized as microtubule-stabilizing (e.g. taxanes) and microtubule-depolymerizing drugs (e.g. vinca alkaloids). Each class shifts the cellular equilibrium that normally exists between soluble tubulin dimers and microtubule polymers, towards microtubules (taxanes) or soluble tubulin (vinca alkaloids). Despite the clinical success of the entire class of microtubule inhibitors, the development of drug resistance – which is the number one cause of cancer mortality in metastatic patients – along with the presence of toxic side effects limits their clinical applicability ^35^. Hence, the discovery of novel microtubule-targeting small molecules could significantly improve cancer therapy by identifying compounds with activity on refractory tumors or compounds with less toxic side effects. To this aim, we further focused our list of high confidence orphan-target predictions to small molecules predicted to target microtubules. To see how our novel predictions related to known microtubule-targeting therapeutics, we created a network of all known and predicted anti-microtubule small molecules with edges representing a predicted shared target interaction (**Figure S7**). Interestingly we found that the 14 known microtubule-targeting agents tended to cluster together based on their distinct mechanism of action. For instance, we observe Paclitaxel clustering with Cabazitaxel and Docetaxel – all known microtubule-stabilizing drugs – while Colchicine clustered with other known microtubule-destabilizing drugs such as Podophyllotoxin. This is especially exciting since it demonstrates the potential for BANDIT to be used not only to identify a specific target for an orphan molecule but to differentiate between different modes of action on the same target.

From our list of top anti-microtubule drug predictions we obtained a set of 24 compounds with varying structures for experimental testing (**Methods, Table S1**). We chose the human breast cancer MDA-MB-231 cells for the validation experiments as microtubule-inhibitors (both stabilizing and destabilizing) are commonly used in the treatment of breast cancer patients. Cells were treated for 6 hours with 1 and 10 μM of each small molecule, and the integrity of the microtubule cytoskeleton (assessed by confocal microscopy following tubulin immunofluorescence), was used as the bio-assay endpoint. Our results showed that 16 of the 24 orphan small molecules exhibited significant effects on microtubules (Figure 3A-F, **Figure S8- 13**), a much higher success rate (67%) than one would expect by chance (p < 2e-16, **Methods**). To more accurately quantify the extent of drug-target engagement, we employed a second biochemical assay quantifying the effect that each small molecule exerted on the equilibrium between microtubule polymers and soluble tubulin, following 6 hours of treatment (**Figure S14**). Our results confirmed and corroborated the microscopy results, further revealing that while several small molecules had maximal microtubule-inhibitory activity at the lowest dose (1μM) (Figure 3C-F), others exhibited a dose-dependent effect on microtubule depolymerization (e.g. compounds #12, #13), further establishing microtubules as their bona-fide target (Figure 3G-I). Taken together, these experiments confirmed the predicted targets and mechanism of action for the majority of the newly identified microtubule inhibitors. While further testing will be needed before these small molecules can be used clinically, these results do demonstrate BANDIT’s target prediction accuracy and how it can be used on compound libraries to identify small molecules acting with a specific mode of action on specific targets, for further investigation.

**Figure 3:**
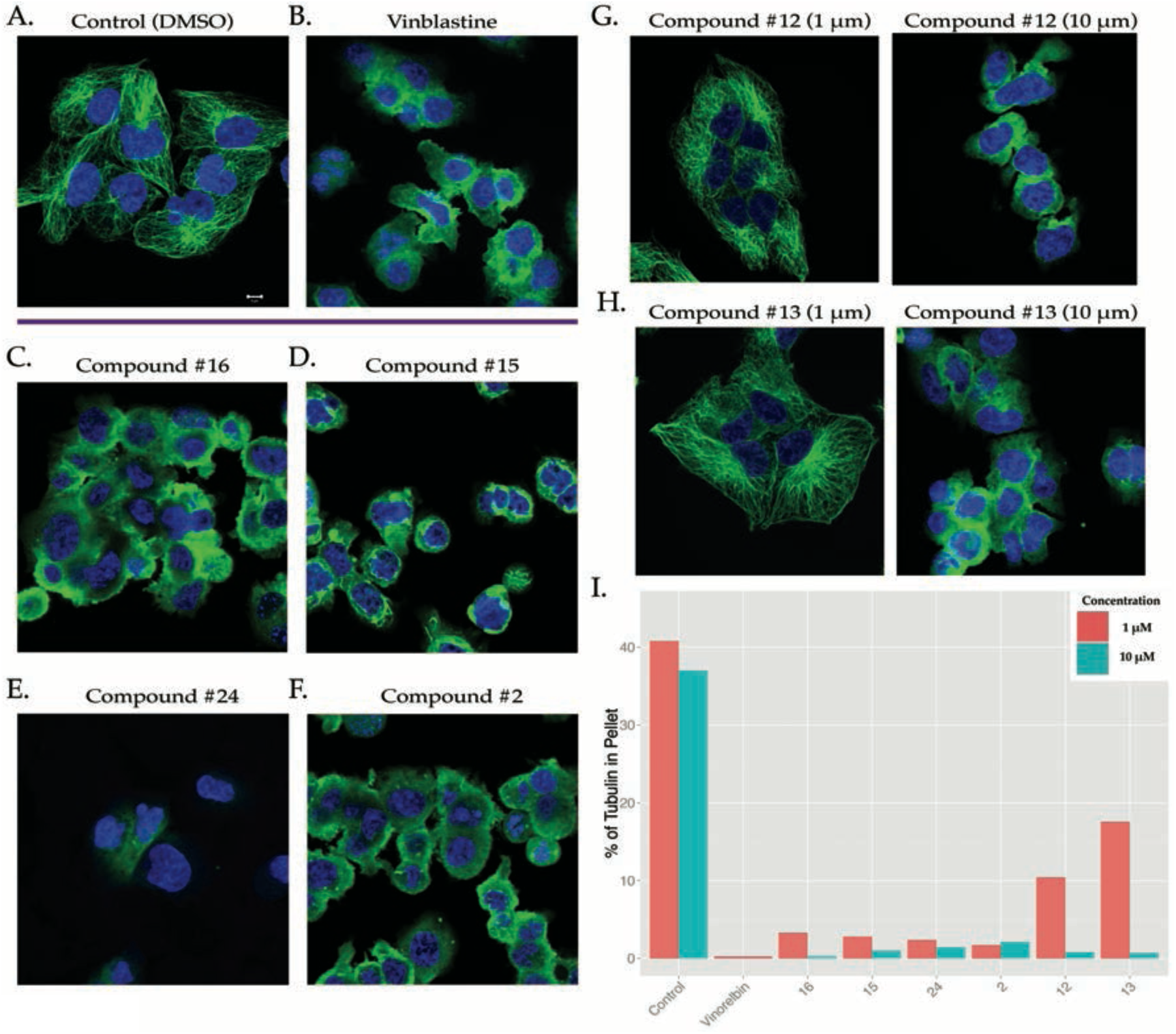
Microtubules are a correct target of the newly identified small molecules – Effect of various compounds (1μM) on the microtubule integrity of MDA-MB-231 cells after 6 hours of treatment. A) Control with DMSO (Scale bar: 5 μm), B) Vinblastine as a positive control, C) Compound #16, D) Compound #15, E) Compound #24 F) Compound #2. G) Dose dependent effect of Compound #12 and H) Compound #13. I) Bar graph showing the % tubulin in the pellet compared to the supernatant (averaged over three independent replicates) for depolymerizing drugs at 1 and 10 μM.

To inform future clinical development for these newly identified microtubule inhibitors, we next tested their activity against drug resistant models. Drug resistance remains one of the most challenging areas in clinical oncology, affecting both broad chemotherapy drugs and targeted-therapies. In the case of microtubule inhibitors, overcoming drug resistance is even more challenging as the mechanisms are often multifactorial. As previously demonstrated, BANDIT can accurately identify a set of structurally diverse small molecules that all bind a common target (in this case microtubules), therefore we next investigated whether any of our newly identified microtubule-depolymerizing small molecules could successfully act on tumors resistant to other known anti-microtubule drugs. Using the 1A9 human ovarian carcinoma cell line – which has previously been used successfully in selecting microtubule-inhibitor resistant clones and for high throughput small molecule screening, ^36-40^ – we created clones resistant to Eribulin mesylate, a microtubule depolymerizing drug that is FDA approved for the treatment of docetaxel-refractory breast cancer patients ^41,42^ (Figure 4A). Interestingly, recent clinical data demonstrated that fewer than 50% of breast cancer patients showed any detectable response after treatment with Eribulin, further highlighting the importance of finding new molecules that share the same validated target but are active against the large population of refractory patients ^43^. Our results, using 72-hr cytotoxicity assays showed that the Eribulin-resistant 1A9 cells (1A9-ERB) were more than 7,000 –fold more resistant to Eribulin than the parental cells and exhibited cross-resistance to all classes of clinically used microtubule-depolymerizing drugs (**Table S2**). To test whether the drug-resistance phenotype was due to impaired drug-target engagement, we treated parental and resistant cells for 6 hr only with 1uM of Eribulin or each of the FDA-approved depolymerizing drugs. Consistent with their drug resistance phenotypes, our results showed lack of drug-induced microtubule depolymerization in 1A9-ERB cells in contrast to the complete depolymerization observed in the microtubule network of drug-sensitive 1A9 parental cells (Figure 4B-C, **Figure S15-16**). These on-target drug efficacy results are in agreement with the lack of antitumor activity revealed by the cytotoxicity data further highlighting the importance of discovering novel small molecules that could act on these refractory tumors. We tested the top 4 performing small molecules (#15, 16, 24, and 2) on the 1A9-ERB cells and found that 3 out of 4 compounds tested, were active against the 1A9-ERB cells and effectively depolymerized microtubules, as evidenced by the diffuse soluble tubulin staining following drug treatment (Figures 4E-F, **Figure S15-16**), in contrast to the fine and intricate microtubule network observed in untreated cells (Figures 4E-A). Compound No 15, which was the most active of the 4 compounds, was tested using cytotoxicity assays and was found to almost completely reverse drug-resistance from 7050-fold observed with Eribulin down to 4-fold (**Table S2**). While further *in vitro* and *in vivo* studies are required for the clinical development of these compounds, these results clearly demonstrate BANDIT’s utility in identifying lead small molecules with potential activity against drug resistance tumor models without the labor-and cost-intensive physical screening of thousands of small molecules. Even though BANDIT is “trained” using a database of drugs with known targets and mechanisms, our results show that it can accurately identify small molecules with distinct modes of action from any known drugs in the training set. This also highlights how BANDIT can pinpoint small molecules from large compound libraries with unique mechanisms that could potentially act on drug resistant cells. Compounds such as these could represent the next generation of clinically developed drugs reducing the need for extensive medicinal chemistry and structure-activity studies, therefore, expediting drug development.

**Figure 4:**
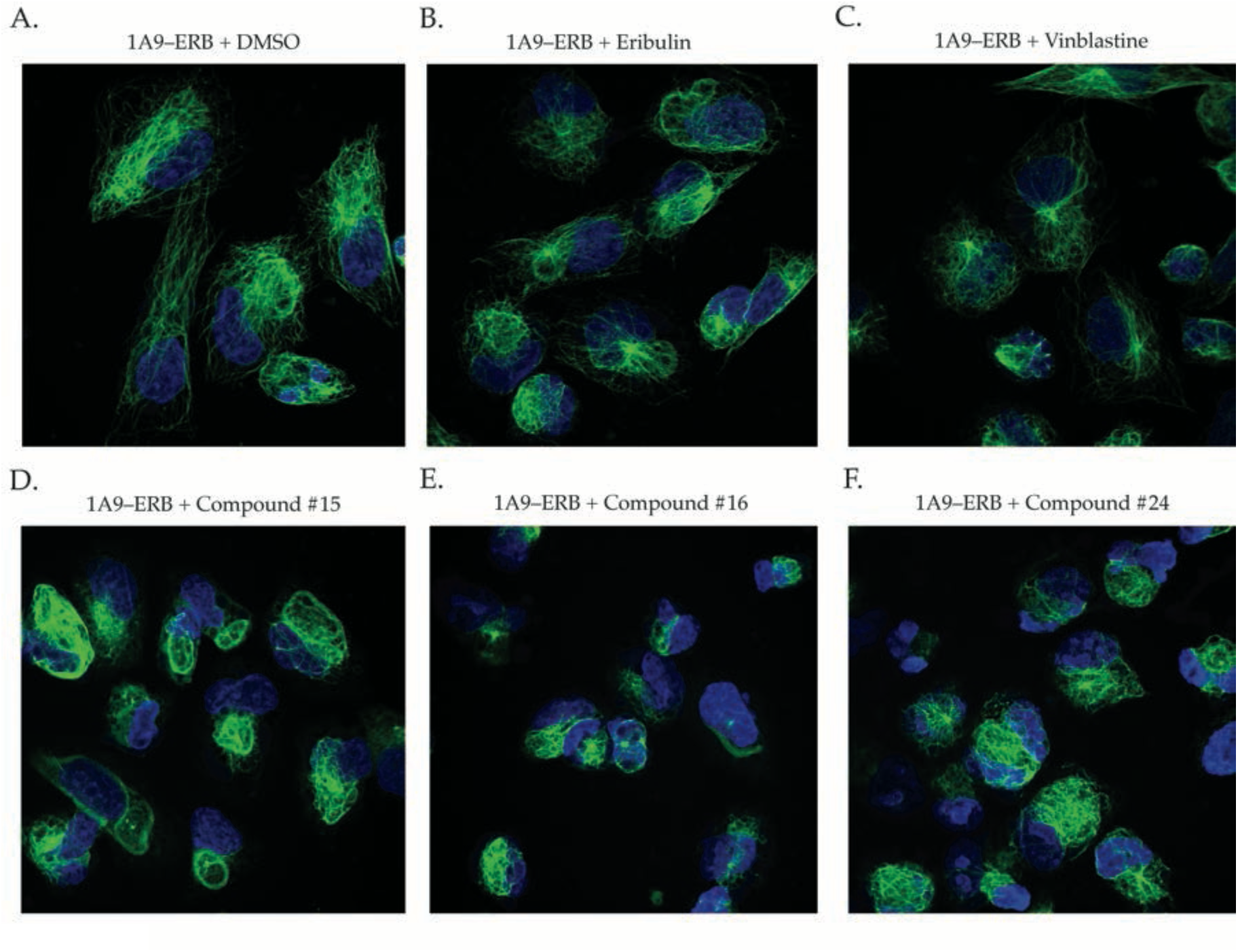
A set of the BANDIT predicted small molecules can act on cells resistant to Eribulin and other microtubule depolymerizing drugs – Effect of various compounds on the microtubule integrity of 1A9-ERB cells after 6 hours of treatment: A) Control with DMSO (Scale bar: 5 μm), 100nM of B) Eribulin and C) Vinblastine, and 1μM of D) Compound #15, E) Compound #16 and F) Compound #24.

### BANDIT Uncovers Selective Antagonism of DRD2 by Anti-Cancer Small Molecule ONC201

Given BANDIT’s demonstrated capability to accurately identify specific targets for orphan small molecules, we next investigated how we could integrate BANDIT directly into the drug development pipeline and test its ability to predict targets for small molecules with promising clinical activity but without a specific target. Therefore we applied BANDIT to ONC201– a small molecule discovered in a phenotypic screen for p53-independent inducers of TRAIL-mediated apoptosis – currently in multiple phase II clinical trials for select advanced cancers. Despite its promising preclinical and early clinical anticancer activity and its reported effects on a few signaling pathways, including Akt/ERK pathway ^44-46^, a bona-fide target for this compound remains elusive.

To identify direct binding targets for ONC201, we used BANDIT to compute likelihood ratios between ONC201 and all drugs with known targets in BANDIT’s database. BANDIT’s top shared target prediction were between ONC201 and Oxiperomide and Thioridazine, both a dopaminergic antagonists previously used the treatment of dyskinesias and schizophrenia respectively ^47-50^. Interestingly, our voting analysis indicated that the most likely targets of ONC201 were dopamine receptors – specifically DRD2 – and adrenergic receptor alpha (Figure 5A), both of which are members of the G-protein coupled receptor (GPCR) superfamily.

**Figure 5:**
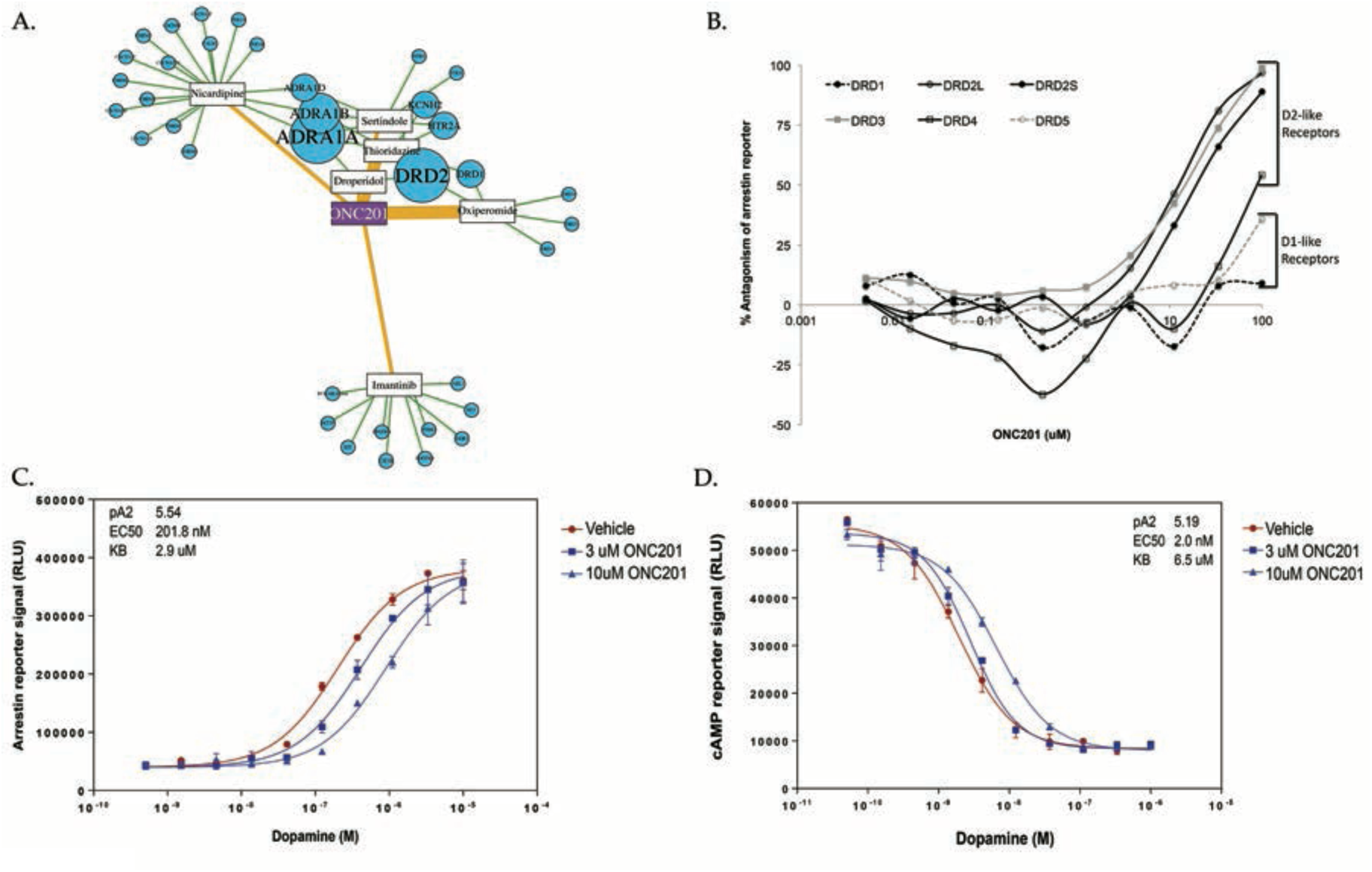
ONC201 is a selective DRD2 antagonist – (A) BANDIT target predictions for ONC201. Connections between ONC201 and known drugs are weighted based on the likelihood ratio and predicted targets are sized based on the prediction strength. (B) Antagonism of ligand-stimulated dopamine receptors by ONC201. C) Schild analysis of DRD2L antagonism by ONC201 using arrestin recruitment or (D) cAMP modulation reporters.

To test these predicted targets we performed in vitro profiling of GPCR activity using a hetereologous reporter assay for arrestin recruitment, which is a hallmark of GPCR activation^51^. Our results indicated that ONC201 selectively antagonized the D2-like (DRD2/3/4L), but not D1- like (DRD1/5L), subfamily of dopamine receptors (Figure 5B; **Figure S17A**), with no observed antagonism of other GPCRs under the evaluated conditions. Among the DRD2 family, ONC201 antagonized both short and long isoforms of DRD2 and DRD3, with weaker potency for DRD4. Further characterization of ONC201-mediated antagonism of arrestin recruitment to DRD2L was assessed by a Gaddam/Schild EC50 shift analysis, which determined a dissociation constant of uM for ONC201 that is equivalent to its effective dose in many human cancer cells (Figure 5C). Confirmatory results were obtained for cAMP modulation in response to ONC201, which is another measure of DRD2L activation (Figure 5D). The ability of dopamine to completely reverse the dose-dependent antagonism of up to 100uM ONC201 suggests direct, competitive antagonism of DRD2L (**Figure S17B-C**). In agreement with the specificity of ONC201 for the target predicted by BANDIT, no significant interactions were identified between ONC201 and nuclear hormone receptors, the kinome, or other drug targets of FDA-approved cancer therapies (**Figure S17D-E**; data not shown). Interestingly, a biologically inactive constitutional isomer of ONC201 ^52^) did not inhibit DRD2L, suggesting that antagonism of this receptor could be linked to its biological activity (**Figure S17F**). In summary, these studies establish that ONC201 selectively antagonizes the D2-like subfamily of dopamine receptors, which is an “unconventional” target for oncology drugs and further demonstrate BANDIT’s ability to act as a tool to advance drug development.

This unexpected discovery on the DRD2L being a direct-binding target for ONC201, has also led to the design and launch of a clinical trial of ONC201 in pheochromocytomas, owing to high levels of DRD2L expression in this rare tumor type. Taken together, these results demonstrate the extreme potential of BANDIT to expedite drug development by using global, novel drug-target engagement predictions in combination with gene expression studies to enable the identification of select patient and indications groups more likely to benefit from a particular drug treatment.

### BANDIT can determine drug mechanisms and can help understand the drug “universe”

Following validation that BANDIT could accurately determine the specific targets for small molecules, we then examined how it could also be used to understand the target binding mechanism, otherwise known as its mechanism of action (MoA). First we used BANDIT to test all known microtubule-targeting drugs, and created a hierarchical cluster based on their TLR outputs (**Methods**). We observed a clean separation between drugs known to destabilize microtubule depolymerizing and polymerizing agents (Figure 6A). A similar MoA-based clustering was observed when we tested all known protein kinase inhibitors, which showed a clear separation between receptor tyrosine kinase inhibitors, serine/threonine kinase inhibitors, and nucleoside analogs (Figure 6B). Overall these results demonstrate that BANDIT can be used to differentiate small molecules based on their specific MoA without additional model training. Combined with the earlier voting algorithm, this demonstrates an efficient pipeline for small molecule target and mechanism identification: first using BANDIT to predict targets for an orphan small molecule, followed by clustering with other drugs known to act on the same target to discern MoA.

**Figure 6:**
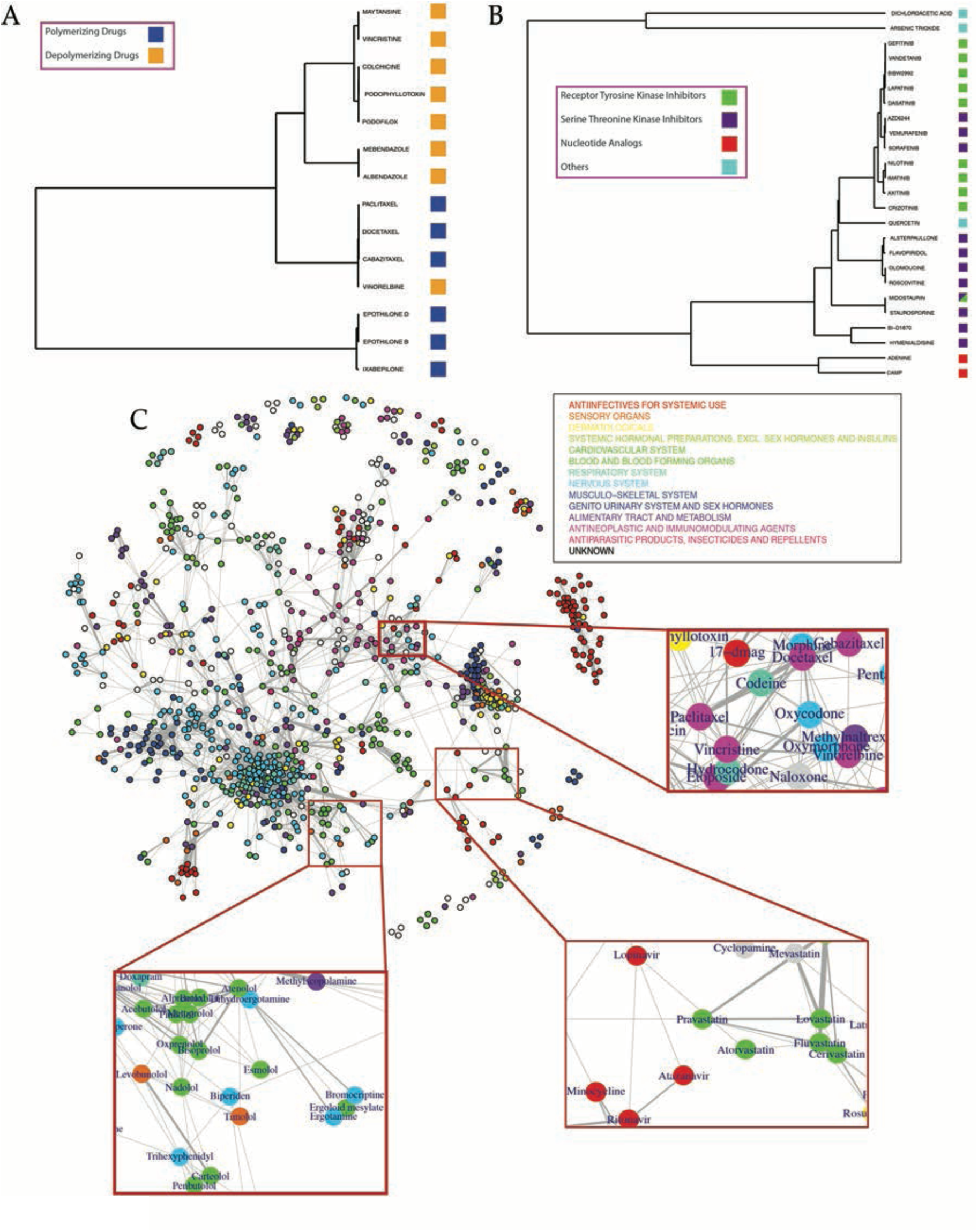
BANDIT can predict specific mechanisms of action and connections between drug classes – A) Hierarchical clustering of drugs known to target microtubules and B) drugs known to target protein kinases. C) Network of drugs based on shared target interactions. Drugs are colored based on their most prevalent ATC code. Three specific clusters corresponding to beta-blockers and Parkinson’s medications, anti-retrovirals and statins, and opioids and anti- microtubule drugs are highlighted.

We next used BANDIT to get an overview of how different classes of drugs, spanning the entire clinical landscape, may be related to one another. Based on the TLR between each drug pair, we constructed a network representative of the drug “universe,” or all known drugs with at least one predicted shared target interaction (Figure 6C). Each drug was classified according to its 1^st^ order Anatomical Therapeutic Chemical (ATC) classification – characteristic of the type and intended use of each drug. As expected, drugs of a similar ATC code cluster together, however we also observed many “unexpected” clusters indicative of drug mechanisms or effect. Interestingly, among all classes of cancer chemotherapeutics, microtubule inhibitors clustered together with camptothecin analogues, for which a dual role as topoisomerase I and tubulin polymerization inhibitors has been previously reported ^53^, but which is not widely acknowledged in clinical oncology. Conversely, we unexpectedly found opioids closely interconnected with microtubule targeting agents; this unanticipated observation is in line with previous reports showing how exposure to microtubule targeting drugs can increase the levels of the opioid receptor in rat cerebellums and that treatment of cardiac myocytes with opioids induces microtubule alterations ^54,55^. This unexploited finding could reveal novel biology linking the opioid receptor signaling pathway with the microtubule cytoskeleton, as well as potentially represent an example of drug repurposing, suggesting novel clinical indications for drugs already FDA-approved. As further proof of the clinical value of the broad universe clustering information revealed by BANDIT, we detected close clustering of known beta-blockers with many Parkinson’s medications, which was especially interesting given that one of the most controversial clinical applications of beta-blockers was to reduce tremors in Parkinson’s patients ^56^. Drug clustering was also strongly indicative of potential side effects, as suggested by the link between antiretroviral medications, which often cause metabolic side effects like hypercholesterolemia, and statins, FDA-approved cholesterol lowering drugs ^57^. Overall we believe this broad universe clustering approach could greatly advance future drug development by “indicating” novel potentially synergistic drug combinations, potentially cumulative side effects, and by assisting in drug repositioning.

## Discussion

One of the strengths of the Bayesian framework is that it can easily accommodate new features, and, as we have observed, we expect that the addition of new data to only improve the overall performance. In addition, as more information becomes available there are many aspects of the current implementation that can be improved. For instance, we can better understand the dependencies between distinct data types and model those within our Bayesian network, and as more information on binding kinetics becomes available, BANDIT could be adapted to better predict on versus off-target effects. As drug development often stops in early clinical studies due to “unanticipated” toxic side effects, BANDIT could help overcome these roadblocks by identifying side effects due to unknown off-target bindings.

In summary, we have developed BANDIT, an integrative Big-Data approach that combines a set of individually weak features into a single reliable and robust predictor of shared-target drug relationships. Not dependent on complex 3D models or large known target cohorts, BANDIT can be used to predict shared target drugs and mechanisms of action for any drug or small molecule (over 50,000 in our database) which differentiates it from other target prediction approaches. By using the top shared-target predictions we can further predict with high accuracy specific targets for a given small molecule and demonstrate how BANDIT can be used to both efficiently discover new drugs with novel mechanisms for specific targets and identify targets for small molecules in the development pipeline – all without tedious, labor-intense and inaccurate drug screening approaches.

Our BANDIT predictions replicated shared-target relationships, individual drug-target relationships, and known mechanisms of action within our test set and replicated results of large-scale experimental screens. Moreover, we experimentally confirmed several of our novel predictions using different bioassays and model systems and demonstrated BANDIT’s capability to efficiently discover novel small molecules, which could be used in refractory tumors. As the development of drug resistance is inevitable in oncology and applicable to both chemotherapy and targeted therapies, BANDIT has the potential to quickly and accurately identify drugs that can potentially overcome resistance and improve patient outcomes. Finally, BANDIT can be used on a broader scale to discern mechanisms of approved drugs, characterize the global drug universe landscape, and explain existing, yet puzzling, clinical phenotypes. That function alone holds tremendous potential for drug repurposing, identification of novel drug combinations, and side effect predictions.

We show herein the potential of BANDIT in expediting drug development, as it spans the entire space ranging from new target-identification and validation to clinical drug development and beyond, by informing repurposing efforts. We expect that BANDIT will help reduce failure rates in the clinic and shorten the time required for drug approval by identifying the right patient population most likely to benefit from a given therapeutic. By allowing researchers to quickly obtain target predictions it could streamline all subsequent drug development efforts and save both time and resources. Furthermore BANDIT could be used to rapidly screen a large database of compounds and efficiently identify any promising therapeutics that could be further evaluated. Overall our results demonstrate that BANDIT is a novel and effective screening and target-prediction platform for drug development and is poised to positively impact current efforts.

## Methods

### Datasets

1. Growth inhibition data: We used publicly available growth inhibition data from the National Cancer Institutes Development Therapeutics Program (NCI-DTP). Each of the NCI60 cell lines were treated with a small molecule and the concentration that caused a 50% decrease in cells was measured. When there were multiple high quality experiments done for the same compound, we averaged the values to obtain a single GI50 value for each small molecule – cell line pair. Contains data on 20,000+ unique compounds. Version 1.6.2 was downloaded from cellminer.com.
2. Gene expression data: All post-treatment gene expression data was downloaded from the Broad Connectivity Map (CMap) project. Fold change data across all cell lines were averaged to obtain a single gene expression signature for each compound. Contains data on 1309 different compounds. Build 02 was downloaded from the Broad CMap Portal.
3. Adverse effects: Side effects (mined from drug package inserts and public information) were downloaded from the SIDER database. Each side effect was classified using the MedDRA (version 16.1) dictionary.
4. Bioassays/Chemical structures: All bioassay results and chemical structures were downloaded from PubChem and organized based on each small molecule’s PubChem Compound Identification (CID).
5. Known Drug Targets: All known drug targets were extracted from the DrugBank database (Version 4.1).

### Calculating similarity scores

1. Growth Inhibition Data: For each pair of drugs we calculated a pearson correlation value across the 60 data points (Figure S1).
2. Gene expression and Chemogenomic Fitness Scores: A pearson correlation was used to measure the degree of similarity for the profiles of two drugs
3. Bioassays: All bioassays were classified as either positive or negative based on the data available in Pubchem. A jaccard index was calculated based on the number of shared “positive” assays between two drugs. We required that each drug pair have been tested in at least one similar assay for a similarity score to be calculated.
4. Chemical Structures: For each drug we extracted the isomeric SMILES and used the atom-pair method ^58^ to calculate the structural similarity between two compounds (Figure S1).
5. Adverse Effects: Using the SIDER2 database ^25^ we extracted the “preferred term” side effects for each drug. A jaccard index was then calculated for the shared side effects for each drug pair.

### Calculating correlations between similarity types

For each pair of similarity scores we separated out drug pairs where both similarity types were measured and plotted the different similarity scores against one another (Figure 1a, Figure S2). We computed the Pearson correlation coefficient (PCC) and the coefficient of determination (R^2^) between each pair of similarity scores. Across all pairs, we observed a low correlation – measured by both the PCC and R^2^. This finding demonstrated that high similarity of one type does not necessarily implied high similarity in another. Furthermore this indicated that each similarity score could be modeled as an independent variable.

### Calculating the Total Likelihood Ratio

For each data type BANDIT calculates a “likelihood ratio” L(s_n_) is defined as the fraction of drug pairs with a shared target (ST pairs) having a given similarity score s_n,_ divided by the fraction of the non-ST pairs with the same similarity score:

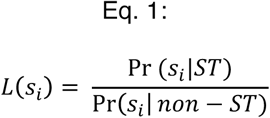

Our previous analysis highlighted the minimal correlation between the similarity types and how data types could be modeled independently under a Naïve Bayes framework. This assumption of independence implies that the joint probability of two drugs sharing a target given a set of similarity scores can be modeled as the product involving individual similarity scores. Therefore the total likelihood ratio L(s) can be expressed as the product of the individual likelihood ratios:

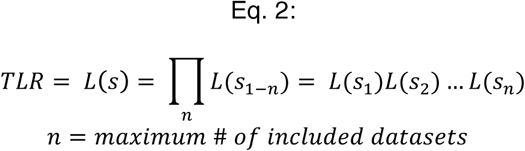

The total likelihood ratio (TLR) is then proportional to the odds of two drugs sharing a given target *n* given sources of information

Overall we decided to use this Bayesian framework for multiple reasons, such as the readily interpretable nature of a likelihood ratio compared to other more complicated machine learning scores and the ability to easily add in new data types as they become available.

### Testing Against Drugs with Known Targets

Drug targets were extracted from DrugBank and drug pairs were classified as a “shared-target” pair if they had at least 1 target in common. We used 5-fold cross validation to split our set of drug pairs into a test and training set containing 20% and 80% of the drug pairs respectively. We sub- sampled the two classes (ST and non-ST drug pairs) and required the ratio of true positives (ST pairs) to true negatives (non-ST pairs) to remain the same as the total set. For each fold we computed TLRs for each drug pair in the test set based on the background probabilities within the training set. Each of the 5 test folds combined at the end to produce an ROC Curve and calculate the AUROC value. We calculated the AUROC value for each individual likelihood ratio from a single data type (Figure S3)

We performed this analysis with the TLR output while varying the number of data types being considered and found a significant increase in the predictive power, measured by the AUROC, as we increased the number of included datasets (Figure 2A). We computed two sets of ROC curves – one where we required drugs have available data in each included data type (our preferred method) and another where we imputed the data type median for each missing data type. We varied the order in which datasets were added and observed a positive relationship between AUROC value and the number of included data types regardless of the addition order. Furthermore we used a KS test to measure how our TLR value could separate out ST and non-ST pairs and saw that in each case our TLR value outperformed any individual variable (Figure S4). We repeated this analysis increasing the minimum number of data types we required a pair of compounds to have and saw the separation steadily improve (D = .44 to .69).

### Replicating Kinase Experimental Screen

We first separated out the kinases in the Peterson et al. database that were classified as BANDIT orphan small molecules – molecules that were in at least two of the considered BANDIT databases and had no known targets. For each orphan kinase inhibitor we used BANDIT to predict shared target drugs. Each known kinase target of the shared target drugs was classified as a potential kinase target of the orphan inhibitor. We then observed that the “percent remaining kinase activity” was significantly lower between the orphan kinase inhibitors and the BANDIT predicted kinases than between the orphan inhibitors and any non-predicted kinases (Wilcoxon Rank Sum Test P = 3.62e–06) (Figure S6).

### Specific Target Voting

For each orphan small molecule we identified all shared target drug predictions, or any drugs with known targets that exceeded a given BANDIT likelihood ratio. For each shared target drug prediction, we compiled all known targets of that given drug and ranked specific protein targets based on how often it appeared as known target in shared drug target predictions. “Votes” for particular protein targets were weighted based on the likelihood ratio of the shared target prediction they originated from. The top voted target for each orphan small molecule that we tested was then predicted to be a novel specific target (Figure 2e).

To test the accuracy, we used leave-one-out cross validation on our test set of drugs with known targets. For each drug we used BANDIT to compare it to all other drugs with known targets and identify the top ranked target for the tested drug. This was repeated for every drug in our test set and we calculated how often the top ranked target was a known target of the drug being tested. We recomputed these accuracies while varying the likelihood ratio cutoff for a drug pair to be considered a shared-target prediction. As expected we observed a steady rise in accuracy as we increased the cutoff value, with the accuracy plateauing at an accuracy level of approximately 90% – revealing that BANDIT’s voting protocol could accurately identify specific targets (Figure 2F).

### Identification of Novel Anti-Microtubule Small Molecules

For each orphan small molecule in BANDIT (defined as a molecule tested in any of the individual databases but without any known targets in DrugBank) we used the BANDIT voting protocol to predict specific protein targets. We required that each orphan small molecule be in at least 3 of BANDIT’s databases, leaving us with a set of ∼15,000 small molecules. To refine our initial list of predictions into a high confidence set, we required a TLR cutoff of 500, that each predicted target appear in the majority of shared target predictions, and that the highest ranked target appear in the top shared target prediction for each orphan molecule. From this list of high confidence predictions we identified a set of small molecules predicted to bind to microtubules.

For each predicted microtubule inhibitor (MTI) we examined how it related to known MTIs using a network approach (Figure S7). We required that each predicted MTI have a TLR greater than 500 with at least two known MTIs. Each edge in our network represents a predicted shared target interaction with the length and width of each corresponding to the strength of the prediction (measured by the TLR value). We used the Fruchterman Reingold projection within the R igraph package. We observed a distinct clustering of known MTIs based on their mechanism of action.

Most of the novel MTIs we predicted were not easily obtained, thus we specifically focused on the subset that we could obtain from the National Cancer Institutes Developmental Therapeutics Program (Table S1).

### Microtubule Imaging/Testing

Human breast MDA-MD-231 cells were cultured in DMEM (obtained from Corning Cellgro) with 10% fetal bovine serum and 1% penicillin and streptomycin. Cells were plated at the density of 90,000 Cells/ml onto 12mm round cover slips in 48 well plates for 24 hours and then treated for 6 hours with small molecules at the given concentrations. Small molecules (obtained from the NCI Drug Bank) were dissolved in DMSO and stored at -20^o^C. Control experiments were done using DMSO and it was less than 0.5% of total media volume. After 6hrs drug treatment media was removed and cells were per-meabilized with 0.5% Triton X-100 and fixed with PHEMO Buffer (3.7% formaldehyde, 0.05% glutaraldehyde, 0.068M Pipes, 0.025M HEPES, 0.015M EGTANa_2_, 0.003M MgCl_2_6H2O and 10% DMSO and adjust pH=6.8) for 10minutes. Fixed cells were washed three times with PBS buffer. Cells were blocked with 10% goat serum at room temperature for 10 minutes. Cells were incubated with monoclonal α-tubulin antibody (clone YL 1/2, obtained from EMD Millipore), for 1hr and washed three times with PBS buffer before incubation with a secondary Alexa Fluor 488 goat anti-mouse antibody (obtained from Invitrogen). Cell chromatin was stained with DAPI for 5min and washed with water three times. Cover slips were mounted and photographed in a RSM 700 microscope for microtubule visualization. DNA was counterstained with DAPI. Images were acquired with Zeiss LSM 700 confocal microscope under a 63×/1.4NA objective (Zeiss, Germany) (Figure 3A-H, Figure S8-S13).

A Fisher’s exact test was used to determine whether the number of observed successes – defined as a predicted microtubule inhibitor showing an effect against microtubules in imaging – was greater than what would be expected by random chance. To determine the background probability we used the number of drugs with known targets in our database that were known to target microtubules (∼ 1%).

### Microtubule Effect Quantification

Following 6hrs treatment, cells (12 well plate) were washed once with warm phosphate-buffered saline. Each well was incubated with 150 μL either with low salts or high salt buffer at 37 ^o^C for 10 minutes. Cell were then scraped and were either lysed in low salt buffer to test for the degree of tubulin polymerization (20 mM Tris–HCl pH 6.8, 1 mM MgCl_2_, 2 mM EGTA, 0.5% NP-40, 1X protease inhibitor cocktail and 0.5% NP-40) or high salt buffer to test for the degree of tubulin depolymerization (0.1M Pipes, 1mM EGTA, 1mM MgSO4, 30% glycerol, 5% DMSO, 1mM DTT, 0.02% NAAzide, 0.125% NP-40, 1mM DTT and 1X protease inhibitor cocktail). Samples were spun at max speed in a tabletop centrifuge for 30 min at room temperature. The supernatant (S) was separated from the pellet (P). The pellet was resuspended in 150 μL 1 × Laemmli buffer and sonicated. Equal volumes of supernatant and pellet samples were loaded onto a 12% gel for a western blot. Tubulin bands were visualized with a DM1α monoclonal antibody (obtained from Sigma-Aldrich). % Tubulin in pellet levels were calculated as the densitometric value of the pellet band divided by the total densitometric value of the pellet and supernatant bands times 100. Three biological repeats were performed (Figure S14).

### Imaging of Treatment Against Resistant Cell Lines

1A9-ERB is a clone of the 1A9 human ovarian carcinoma cell line resistant to the effects of Eribulin mesylate. It was prepared by exposing 1A9 cells to 1ng/ml Eribulin (obtained from Eisai pharmaceuticals) in the presence of 10ug/ml verapamil (obtained from Acros Organics), a Pgp antagonist. The cells were maintained in the 0.5ng/ml eribulin and 10ug/ml verapamil concentrations. Cells were removed from this drug solution 3 days prior to any future experimentation. Additional treatment and imaging was done using the same protocols as described earlier (Figure S15-S17).

### Characterization of ONC201-DRD2 Interaction

ONC201 dihydrochloride was obtained from Oncoceutics. Kinase inhibition assays for the kinome were performed as previously described ^59^. GPCR arrestin recruitment and cAMP modulation reporter assays were performed as previously described ^60^. PathHunter™ (DiscoveRx) beta- arrestin cells expressing one of several GPCR targets were plated onto 384-well white solid bottom assay plates (Corning 3570) at 5000 cells per well in a 20 *μ*L volume in the appropriate cell plating reagent. Cells were incubated at 37 °C, 5% CO_2_ for 18-24 h. Samples were prepared in buffer containing 0.05% fatty-acid free BSA (Sigma). For agonist mode tests, samples (5 *μ*L) were added to pre-plated cells and incubated for 90 minutes at 37 °C, 5% CO_2_. For antagonist mode tests, samples (5 *μ*L) were added to pre-plated cells and incubated for 30 minutes at 37 °C, 5% CO_2_ followed by addition of EC80 agonist (5 *μ*L) for 90 minutes at 37 °C, 5% CO_2_. For Schild analysis, samples (5 *μ*L) were added to pre-plated cells and incubated for 30 minutes at 37 °C, 5% CO_2_ followed by addition of serially dliuted agonist (5 *μ*L) for 90 minutes at 37 °C, 5% CO_2_. Control wells defining the maximal and minimal response for each assay mode were tested in parallel. Arrestin recruitment was measured by addition of 15 *μ*L PathHunter Detection reagent and incubated for 1-2 h at room temperature and read on a Perkin Elmer Envision Plate Reader. For agonist and antagonist tests, data was normalized for percent efficacy using the appropriate controls and fitted to a sigmoidal dose-response (variable slope), Y=Bottom + (Top- Bottom)/(1+10^((LogEC50-X)*HillSlope)), where X is the log concentration of compound.

For Schild analysis, data was normalized for percent efficacy using the appropriate controls and fitted to a Gaddum/Schild EC50 shift using global fitting, where Y=Bottom + (Top- Bottom)/(1+10^((LogEC-X)*HillSlope)), Antag=1+(B/(10^(-1*pA2)))^SchildSlope and LogEC=Log(EC50*Antag). EC50 / IC50 analysis was performed in CBIS data analysis suite (Cheminnovation) and Schild analysis performed in GraphPad Prism 6.0.5 (Figure 5, Figure S17).

The kinase assay and nuclear hormone receptor profiling (S16) were performed as previously described by Reaction Biology Corp and DiscoverX respectively ^61-63^.

### Drug Mechanism Clustering

For each drug pair we converted the TLR between them into a distance metric used to estimate “closeness” between any two drugs:

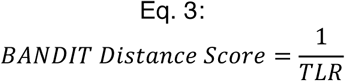

We next separated all drugs know to target microtubules that were in at least 3 of BANDIT’s dataset. With the BANDIT distance metric as an input we created a hierarchical cluster of all known MTIs using the hclust R method with an “average” based clustering method. Known MTIs were labeled based on whether they were known to polymerize or depolymerize microtubules, and we observed a distinct separation based on the mechanism of action (MoA). We repeated this clustering while removing drug structures from our likelihood calculations and continued to see a MoA-based separation (Figure S18). This revealed that BANDIT’s clustering approach is not dependent on any single data type, and that observed results are due to BANDIT’s integrative approach. This analysis was then repeated using similar conditions for known protein kinases.

### Drug “Universe” Clustering

Using the same protocol as was used to create the MTI network, we created a network of all drugs with known targets with each edge representing a predicted shared target interaction and the edge weight corresponding to the strength of the interaction. Using the KEGG drug database^64^ and DrugBank^27^ we annotated each drug based on its most prevalent ATC code and colored each drug accordingly. We specifically isolated out 3 clusters representing: 1) beta-blockers with Parkinson’s medications, 2) antiretrovirals and statins and 3) opioids and microtubule inhbitors.

To get a better understanding of how orphan small molecules fit into this drug “universe” we computed the distance between every pair of small molecules and used multi-dimensional scaling to visualize the overall structure (Figure S19). We used the same distance metric as described in the mechanism of action clustering section to create a distance matrix between all small molecules (known drugs and orphan) and used the R cmdscale package for the multi-dimensional scaling. We noticed a definite structure with known drugs tightly clustering around each other, while orphan molecules had a more diffuse organization. One explanation for this structure is that drugs with known targets are more likely to be used to treat patients and thus may have similar effects due to safety precautions, whereas orphan molecules which have not gone through clinical trials and FDA approval are more likely to have a wide variety of effects and characteristics.

## References

1 Cuatrecasas, P. Drug discovery in jeopardy. The Journal of clinical investigation 116, 2837–2842, doi:10.1172/JCI29999 (2006).

2 Chan, J. N., Nislow, C. & Emili, A. Recent advances and method development for drug target identification. Trends in pharmacological sciences 31, 82–88, doi:10.1016/j.tips.2009.11.002 (2010).

3 Weigelt, J. The case for open-access chemical biology. A strategy for precompetitive medicinal chemistry to promote drug discovery. EMBO reports 10, 941–945, doi:10.1038/embor.2009.193 (2009).

4 Williams, M. Target validation. Current opinion in pharmacology 3, 571–577 (2003).

5 Dearden, J. C. In silico prediction of drug toxicity. Journal of computer-aided molecular design 17, 119–127 (2003).

6 Butina, D., Segall, M. D. & Frankcombe, K. Predicting ADME properties in silico: methods and models. Drug Discov Today 7, S83–88 (2002).

7 Nantasenamat, C., Isarankura-Na-Ayudhya, C. & Prachayasittikul, V. Advances in computational methods to predict the biological activity of compounds. Expert Opin Drug Discov 5, 633–654, doi:10.1517/17460441.2010.492827 (2010).

8 Li, H. et al. TarFisDock: a web server for identifying drug targets with docking approach. Nucleic acids research 34, W219–224, doi:10.1093/nar/gkl114 (2006).

9 Rarey, M., Kramer, B., Lengauer, T. & Klebe, G. A fast flexible docking method using an incremental construction algorithm. Journal of molecular biology 261, 470–489, doi:10.1006/jmbi.1996.0477 (1996).

10 Wang, K. et al. Prediction of drug-target interactions for drug repositioning only based on genomic expression similarity. PLoS computational biology 9, e1003315, doi:10.1371/journal.pcbi.1003315 (2013).

11 Lamb, J. The Connectivity Map: a new tool for biomedical research. Nature reviews. Cancer 7, 54–60, doi:10.1038/nrc2044 (2007).

12 Lamb, J. et al. The Connectivity Map: using gene-expression signatures to connect small molecules, genes, and disease. Science 313, 1929–1935, doi:10.1126/science.1132939 (2006).

13 Campillos, M., Kuhn, M., Gavin, A. C., Jensen, L. J. & Bork, P. Drug target identification using side-effect similarity. Science 321, 263–266, doi:10.1126/science.1158140 (2008).

14 Grundmark, B., Holmberg, L., Garmo, H. & Zethelius, B. Reducing the noise in signal detection of adverse drug reactions by standardizing the background: a pilot study on analyses of proportional reporting ratios-by-therapeutic area. Eur J Clin Pharmacol 70, 627–635, doi:10.1007/s00228-014-1658-1 (2014).

15 Shang, N., Xu, H., Rindflesch, T. C. & Cohen, T. Identifying plausible adverse drug reactions using knowledge extracted from the literature. J Biomed Inform 52, 293–310, doi:10.1016/j.jbi.2014.07.011 (2014).

16 Fortney, K. et al. Prioritizing therapeutics for lung cancer: an integrative meta-analysis of cancer gene signatures and chemogenomic data. PLoS computational biology 11, e1004068, doi:10.1371/journal.pcbi.1004068 (2015).

17 Ma’ayan, A. et al. Lean Big Data integration in systems biology and systems pharmacology. Trends in pharmacological sciences 35, 450–460, doi:10.1016/j.tips.2014.07.001 (2014).

18 Wang, Z., Clark, N. R. & Ma’ayan, A. Drug-induced adverse events prediction with the LINCS L1000 data. Bioinformatics 32, 2338–2345, doi:10.1093/bioinformatics/btw168 (2016).

19 Perlman, L., Gottlieb, A., Atias, N., Ruppin, E. & Sharan, R. Combining drug and gene similarity measures for drug-target elucidation. Journal of computational biology: a journal of computational molecular cell biology 18, 133–145, doi:10.1089/cmb.2010.0213 (2011).

20 Fakhraei, S., Huang, B., Raschid, L. & Getoor, L. Network-Based Drug-Target Interaction Prediction with Probabilistic Soft Logic. IEEE/ACM transactions on computational biology and bioinformatics / IEEE, ACM 11, 775–787, doi:10.1109/TCBB.2014.2325031 (2014).

21 Chen, X. et al. Drug-target interaction prediction: databases, web servers and computational models. Brief Bioinform 17, 696–712, doi:10.1093/bib/bbv066 (2016).

22 Shoemaker, R. H. The NCI60 human tumour cell line anticancer drug screen. Nature reviews. Cancer 6, 813–823, doi:10.1038/nrc1951 (2006).

23 Li, Q., Cheng, T., Wang, Y. & Bryant, S. H. PubChem as a public resource for drug discovery. Drug Discov Today 15, 1052–1057, doi:10.1016/j.drudis.2010.10.003 (2010).

24 Chen, B. & Wild, D. J. PubChem BioAssays as a data source for predictive models. Journal of molecular graphics & modelling 28, 420–426, doi:10.1016/j.jmgm.2009.10.001 (2010).

25 Kuhn, M., Campillos, M., Letunic, I., Jensen, L. J. & Bork, P. A side effect resource to capture phenotypic effects of drugs. Molecular systems biology 6, 343, doi:10.1038/msb.2009.98 (2010).

26 Law, V. et al. DrugBank 4.0: shedding new light on drug metabolism. Nucleic Acids Res 42, D1091–1097, doi:10.1093/nar/gkt1068 (2014).

27 Wishart, D. S. et al. DrugBank: a knowledgebase for drugs, drug actions and drug targets. Nucleic acids research 36, D901–906, doi:10.1093/nar/gkm958 (2008).

28 Yamanishi, Y., Kotera, M., Kanehisa, M. & Goto, S. Drug-target interaction prediction from chemical, genomic and pharmacological data in an integrated framework. Bioinformatics 26, i246–254, doi:10.1093/bioinformatics/btq176 (2010).

29 Hizukuri, Y., Sawada, R. & Yamanishi, Y. Predicting target proteins for drug candidate compounds based on drug-induced gene expression data in a chemical structure-independent manner. BMC Med Genomics 8, 82, doi:10.1186/s12920-015-0158-1 (2015).

30 Anastassiadis, T., Deacon, S. W., Devarajan, K., Ma, H. & Peterson, J. R. Comprehensive assay of kinase catalytic activity reveals features of kinase inhibitor selectivity. Nat Biotechnol 29, 1039–1045, doi:10.1038/nbt.2017 (2011).

31 Jordan, M. A. & Wilson, L. Microtubules as a target for anticancer drugs. Nature reviews. Cancer 4, 253–265 (2004).

32 Jordan, M. A. & Wilson, L. Microtubules and actin filaments: dynamic targets for cancer chemotherapy. Curr Opin Cell Biol 10, 123–130 (1998).

33 Giannakakou, P., Sackett, D. & Fojo, T. Tubulin/microtubules: still a promising target for new chemotherapeutic agents. J Natl Cancer Inst 92, 182–183 (2000).

34 Jordan, A., Hadfield, J. A., Lawrence, N. J. & McGown, A. T. Tubulin as a target for anticancer drugs: Agents which interact with the mitotic spindle. Medicinal Research Reviews 18, 259–296, doi:10.1002/(SICI)1098- 1128(199807)18:4259::AID-MED33.0.CO;2-U (1998).

35 Mukhtar, E., Adhami, V. M. & Mukhtar, H. Targeting microtubules by natural agents for cancer therapy. Molecular cancer therapeutics 13, 275–284, doi:10.1158/1535-7163.MCT-13-0791 (2014).

36 Giannakakou, P. et al. Paclitaxel-resistant human ovarian cancer cells have mutant beta-tubulins that exhibit impaired paclitaxel-driven polymerization. The Journal of biological chemistry 272, 17118–17125 (1997).

37 Nicolaou, K. C. et al. Synthesis of epothilones A and B in solid and solution phase. Nature 387, 268–272, doi:10.1038/387268a0 (1997).

38 Giannakakou, P. et al. A common pharmacophore for epothilone and taxanes: molecular basis for drug resistance conferred by tubulin mutations in human cancer cells. Proceedings of the National Academy of Sciences of the United States of America 97, 2904–2909, doi:10.1073/pnas.040546297 (2000).

39 Nicolaou, K. C. et al. Chemical synthesis and biological evaluation of cis- and trans-12,13-cyclopropyl and 12,13-cyclobutyl epothilones and related pyridine side chain analogues. J Am Chem Soc 123, 9313–9323 (2001).

40 Nicolaou, K. C. et al. Design, synthesis, and biological properties of highly potent epothilone B analogues. Angew Chem Int Ed Engl 42, 3515–3520, doi:10.1002/anie.200351819 (2003).

41 O’Rourke, B., Yang, C. P., Sharp, D. & Horwitz, S. B. Eribulin disrupts EB1- microtubule plus-tip complex formation. Cell Cycle 13, 3218–3221, doi:10.4161/15384101.2014.950143 (2014).

42 Dybdal-Hargreaves, N. F., Risinger, A. L. & Mooberry, S. L. Eribulin mesylate: mechanism of action of a unique microtubule-targeting agent. Clin Cancer Res 21, 2445–2452, doi:10.1158/1078-0432.CCR-14-3252 (2015).

43 Gamucci, T. et al. Eribulin mesylate in pretreated breast cancer patients: a multicenter retrospective observational study. J Cancer 5, 320–327, doi:10.7150/jca.8748 (2014).

44 Allen, J. E. et al. Dual Inactivation of Akt and ERK by TIC10 Signals Foxo3a Nuclear Translocation, TRAIL Gene Induction, and Potent Antitumor Effects. Science translational medicine 5, 171ra117–171ra117, doi:10.1126/scitranslmed.3004828 (2013).

45 J. Ishizawa et al. ONC201 Induces p53-independent Apoptosis in Hematological Malignancies and Leukemic Stem/Progenitor Cells through ER Stress Response. Science Signaling (2015 (in press)).

46 Kline CL et al. Anti-cancer agent ONC201 activates early ATF4/DR5 upregulation, and cell death associated with XIAP inhibition. Science Signaling (2015 (in press)).

47 Bedard, P., Parkes, J. D. & Marsden, C. D. Effect of new dopamine-blocking agent (oxiperomide) on drug-induced dyskinesias in Parkinson’s disease and spontaneous dyskinesias. Br Med J 1, 954–956 (1978).

48 Casey, D. E. & Gerlach, J. Oxiperomide in tardive dyskinesia. J Neurol Neurosurg Psychiatry 43, 264–267 (1980).

49 Casey, D. E. & Gerlach, J. Sulpiride and oxiperomide in tardive dyskinesia. Trans Am Neurol Assoc 104, 210–211 (1979).

50 Meltzer, H. Y., Sachar, E. J. & Frantz, A. G. Dopamine antagonism by thioridazine in schizophrenia. Biol Psychiatry 10, 53–57 (1975).

51 Zhang, R. & Xie, X. Tools for GPCR drug discovery. Acta Pharmacol Sin 33, 372–384, doi:10.1038/aps.2011.173 (2012).

52 Wagner, J. et al. The angular structure of ONC201, a TRAIL pathway-inducing compound, determines its potent anti-cancer activity. Oncotarget 5, 12728– 12737 (2014).

53 Chang, J. Y. et al. Dual inhibition of topoisomerase I and tubulin polymerization by BPR0Y007, a novel cytotoxic agent. Biochem Pharmacol 65, 2009–2019 (2003).

54 Devillard, L. et al. Opioid-induced protection of cardiac myocytes from ischemic injury: Involvement of microtubules. Journal of Molecular and Cellular Cardiology 42, S193–S194, doi:10.1016/j.yjmcc.2007.03.588.

55 Borsodi, A. & Toth, G. Microtubule disassembly increases the number of opioid receptor binding sites in rat cerebrum membranes. Neuropeptides 8, 51–54 (1986).

56 Crosby, N. J., Deane, K. H. & Clarke, C. E. Beta-blocker therapy for tremor in Parkinson’s disease. Cochrane Database Syst Rev, CD003361, doi:10.1002/14651858.CD003361 (2003).

57 Carr, A. & Cooper, D. A. Adverse effects of antiretroviral therapy. Lancet 356, 1423–1430, doi:10.1016/S0140-6736(00)02854-3 (2000).

58 Carhart, R. E., Smith, D. H. & Venkataraghavan, R. Atom pairs as molecular features in structure-activity studies: definition and applications. Journal of Chemical Information and Computer Sciences 25, 64–73, doi:10.1021/ci00046a002 (1985).

59 Anastassiadis, T., Deacon, S. W., Devarajan, K., Ma, H. & Peterson, J. R. Comprehensive assay of kinase catalytic activity reveals features of kinase inhibitor selectivity. Nat Biotech 29, 1039–1045, doi: http://www.nature.com/nbt/journal/v29/n11/abs/nbt.2017.html-supplementary-information (2011).

60 McGuinness, D. et al. Characterizing Cannabinoid CB 2 Receptor Ligands Using DiscoveRx PathHunter(tm) ß-Arrestin Assay. Journal of Biomolecular Screening 14, 49–58, doi:10.1177/1087057108327329 (2009).

61 Patel, A. et al. A combination of ultrahigh throughput PathHunter and cytokine secretion assays to identify glucocorticoid receptor agonists. Anal Biochem 385, 286–292, doi:10.1016/j.ab.2008.11.005 (2009).

62 Corp, R. B. Reaction Biology Corp Kinase Assay Protocol, <http://www.reactionbiology.com/webapps/site/Kinase_Assay_Protocol.aspx> (2017).

63 DiscoverX. PathHunter Nuclear Translocation Assays, <https://www.discoverx.com/technologies-platforms/enzyme-fragment-complementation-technology/cell-based-efc-assays/protein-translocation/nuclear-translocation-assays> (2017).

64 Kanehisa, M. & Goto, S. KEGG: Kyoto Encyclopedia of Genes and Genomes. Nucleic acids research 28, 27–30, doi:DOI 10.1093/nar/28.1.27 (2000).

